# LOSS OF CLDN5 -AND INCREASE IN IRF7- IN THE HIPPOCAMPUS AND CEREBRAL CORTEX OF DIABETIC MICE AT THE EARLY SYMPTOMATIC STAGE

**DOI:** 10.1101/2023.11.02.565280

**Authors:** Marta Carús-Cadavieco, Sandra González de la Fuente, Inés Berenguer, Miguel A. Serrano-Lope, Begoña Aguado, Ernest Palomer, Francesc Guix, Carlos G. Dotti

## Abstract

Analyzing changes in gene expression within specific brain regions of individuals with Type 2 Diabetes (T2DM) who do not exhibit significant cognitive deficits can yield valuable insights into the mechanisms that may underlie the progression toward a more severe phenotype, for example as when individuals age. Here, we present evidence that adult mice with long-term type 2 diabetes mellitus (T2DM) and minor cognitive deficits display alterations in the expression of 27 genes in the cerebral cortex and 16 genes in the hippocampus compared to non-T2DM mice. Only six of these genes undergo the same type of change both in the cortex and hippocampus: Interferon regulatory factor 7 (Irf7), Hypoxia-inducible factor 3 alpha (*Hif*-3α), period circadian clock 2 (*Per2*), xanthine dehydrogenase (*Xdh*), and Transforming growth factor β-stimulated clone 22/TSC22 (*Tscd3*) are all upregulated, while Claudin-5 (*Cldn5*) is downregulated. At the protein level, Claudin5 and IRF7 showed equivalente changes: downregulation of CLDN5 and upregulation of IRF7. These results suggest that cognitive deficits linked to chronic T2DM may stem from compromised blood-brain barrier integrity and an abnormal inflammatory response in the early stages of the disease. This underscores the potential for therapeutic interventions targeting CLDN5 and IRF7.

## 1. Background

Numerous clinical studies have unveiled that individuals with long-term T2DM may exhibit mild cognitive impairments, especially in areas such as verbal memory, information processing speed, attention, and task execution [1,2]. In line with these observations, we have recently shown that adult mice with long-term T2DM display only minor learning impairments [3]. Acknowledging that the prolonged influence of T2DM could result in more profound cognitive deficits, and possibly even initiate the emergence of neurodegeneration [1,4,5], we have here analysed the pattern of gene expression of the cortex and hippocampus of these adult, chronically T2DM, mice.

## 2. Methods

Mice between 7 and 9 months of age were subjected to HFD for 14 weeks, at which time an intraperitoneal injection of STZ (40mg/kg) per day was performed for 5 days [3,6]. Although STZ is a compound commonly used as an experimental model of type 1 diabetes (since this antibiotic induces the destruction of pancreatic ß cells leading to hypoinsulinemia and hyperglucemia), its use at multiple low doses and in combination with HFD allows it to mimic the final stages of T2DM [6]. After the STZ treatment, HFD was continued for another 6 weeks (age of the mice at the end of the experiment was 12-14 months). Behavioral tests, immunoblotting, antibodies, RNA-sequencing and data analysis, transcript enrichment analysis and statistical analysis have been described in the previous publication [3] and are also summarized in the legends of figures and as supplementary information.

## 3. Results

In a recent publication we showed that wild type mice subjected to the HFD and STZ experimental setup develop classical signs of metabolic syndrome, including weight gain, hyperglycaemia and impaired response to the glucose and insulin tolerance tests [3]. In the same work, subjecting these mice to various cognitive assessments, including the Y maze, open field, novel object recognition, and Barnes maze, revealed -minor-deficits in the Barnes and the Y maze [3]. To gain mechanistic insights on how T2DM might be leading to these brain function changes, we carried out a transcriptomic analysis of cortical and hippocampal samples from control and the same T2DM mice used for the behavioral studies. Total RNA from the cortex and hippocampus of 3 control and 3 T2DM mice was extracted and after poly(A)+ selection and RNA sequencing was carried out using the Illumina methodology as described previously [3]. Tables 1 and 2 summarize the RNA-Seq datasets generated for this study. The Tables illustrate that T2DM produces significant changes in the expression of 27 genes in the cerebral cortex (Table 1) and 16 in the hippocampus (Table 2). To depict gene expression differences, a log2 fold change (log2FC) bar plot of differentially expressed genes (DEGs) from both cortex and hippocampus comparatives, was crafted. This analysis revealed changes in expression levels of 28 genes in the cerebral cortex and 15 genes in the hippocampus. Fig. 1A illustrates in a graphical manner the quantitative differences of T2DM on gene expression (including both cortex and hippocampus DEGs): 24 genes are upregulated and 19 downregulated. Next, an overrepresentation analysis (ORA) [7] using the ClusterProfiler R package [8] (see Supplementary materials) was conducted to determine whether known biological functions or processes are over-represented in the experimentally-derived gene list. The gene ontology (GO) plot in Figure 1B offers insights into the pathways that could be most influenced by the up and downregulation of these genes: inflammation and immunity (e.g. *Irf7, Irf9, Ifit1, GM19439, Ly6C*), cellular rhythmicity (e.g. *Ciart, Per2, Dbp, Tef)*, metabolism (*AMPK, TSC22, AdipoR2, Mt2, Fkbp5, AsgR1, Cyp26b*), maintenance of the blood-brain barrier/vascular integrity (e.g. *Claudin 5, Cav1, MMP14, Sdc4)*, oxidative metabolism/response to hypoxia (e.g. *Gpt2, Xdh, Hif3a, TSC22, Mt2*), gene expression (*Xlr4a*), cell-cell interaction/morphogenesis (*Xlr4a, Hspg2, Sdc4, Zic4, Bmp6, Sned1, Col1a*), and neurotransmission/synaptic plasticity (e.g. *Xlr4a, CamK1, Slc6a13*) [9–13]. Considering that any of the DEGs in T2DM-induced mice could favour cognitive disorders and that common changes in the cortex and hippocampus would have stronger functional implications, we next analyzed if there were genes that changed in the same fashion in the two structures. Our results showed that 6 genes undergo similar changes in expression pattern in the two brain structures: while *Hif*-3α, IrfF7, Per2, Xdh, and Tsc22/Tscd3 are upregulated, Cldn 5 is downregulated (Fig.2A). Given that there is not always a correlation between mRNA and protein levels (i.e. translation efficiency, protein degradation, protein stability, feedback loops and compensatory mechanisms), we next analyzed the protein levels of these 6 transcripts. Figure 2B-C shows that T2DM induces changes in CLDN5 and IRF7 at the protein level in the hippocampus that parallel the mRNA changes: reduced CLDN5 and increased IRF7.

**TABLE 1:**
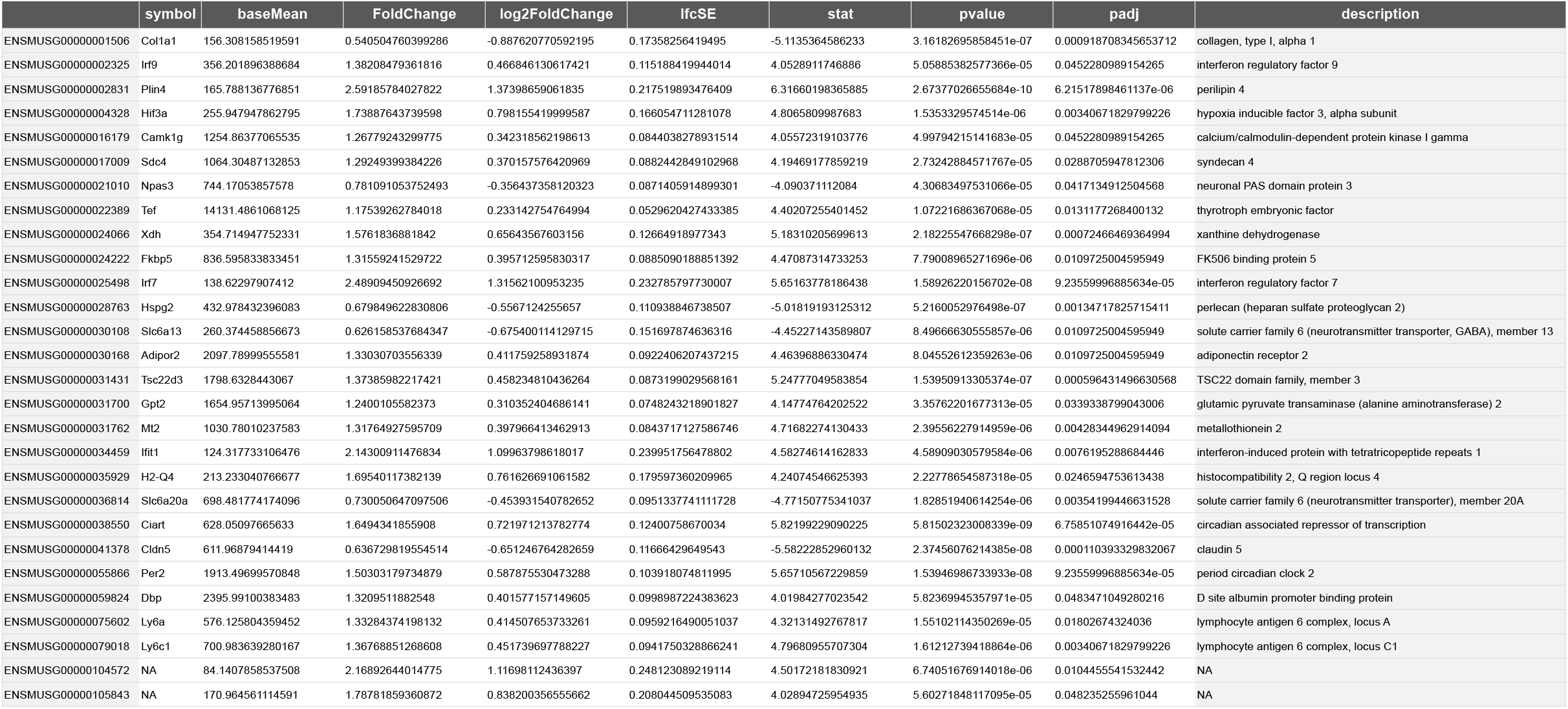
Gene Expression changes in the Cortex.

**TABLE 2:** Gene Expression changes in the Hippocampus.

|  | symbol | baseMean | FoldChange | log2FoldChange | lfcSE | stat | pvalue | padj | description |
| --- | --- | --- | --- | --- | --- | --- | --- | --- | --- |
| ENSMUSG000000000957 | Mmp14 | 552.306524199356 | 0.621477405415459 | -0.6862261536299 | 0.165606113822399 | -4.14372475623593 | 3.41709845359327e-05 | 0.0445849120371853 | matrix metallopeptidase 14 (membrane-inserted) |
| ENSMUSG000000004044 | Cavin1 | 609.226522377664 | 0.619365789094502 | -0.691136397484694 | 0.164668268872255 | -4.19714376192815 | 2.7030236163915e-05 | 0.0420603493640707 | caveolae associated 1 |
| ENSMUSG000000004328 | Hif3a | 263.067022966399 | 2.00147952385081 | 1.00106685630002 | 0.180421284149632 | 5.54849645937443 | 2.88136718896591e-08 | 0.000280774825728783 | hypoxia inducible factor 3, alpha subunit |
| ENSMUSG000000020884 | Asgr1 | 26.8581133329989 | 0.1097190482604 | -3.18811408195198 | 0.75691115596043 | -4.21200567179729 | 2.53113187278517e-05 | 0.0420603493640707 | asialoglycoprotein receptor 1 |
| ENSMUSG000000024066 | Xdh | 336.21373210551 | 1.54376232807968 | 0.626450657726244 | 0.151215857393462 | 4.14275770097461 | 3.43154436121801e-05 | 0.0445849120371853 | xanthine dehydrogenase |
| ENSMUSG000000025498 | Irf7 | 134.960155998884 | 2.37273052908707 | 1.2465482630161 | 0.288831198199165 | 4.3158366228725 | 1.58999548546372e-05 | 0.0420603493640707 | interferon regulatory factor 7 |
| ENSMUSG000000031431 | Tsc22d3 | 1841.04582691217 | 1.41999052082298 | 0.505881299020575 | 0.119476586305837 | 4.23414590809965 | 2.29421959148577e-05 | 0.0420603493640707 | TSC22 domain family, member 3 |
| ENSMUSG000000036972 | Zic4 | 183.703350061377 | 0.289564828996892 | -1.78804171385155 | 0.348113099814262 | -5.13638158060002 | 2.80078695197622e-07 | 0.00109169073814129 | zinc finger protein of the cerebellum 4 |
| ENSMUSG000000038205 | Prkab2 | 2480.99798051789 | 1.27311387932047 | 0.348361473271261 | 0.0814070436372269 | 4.27925468984796 | 1.87520186898725e-05 | 0.0420603493640707 | protein kinase, AMP-activated, beta 2 non-catalytic subunit |
| ENSMUSG000000039004 | Bmp6 | 381.798435688192 | 0.406603242554175 | -1.29830637454443 | 0.309954706232084 | -4.18869708521959 | 2.80560594044291e-05 | 0.0420603493640707 | bone morphogenetic protein 6 |
| ENSMUSG000000041378 | Cldn5 | 760.496801886941 | 0.591236783133838 | -0.75819206695608 | 0.140681814544885 | -5.38941063142298 | 7.06891211763785e-08 | 0.000459220094202147 | claudin 5 |
| ENSMUSG000000047793 | Sned1 | 954.621087281502 | 0.515677928856565 | -0.955457795734936 | 0.122121556042922 | -7.82382592143784 | 5.12416535574678e-15 | 9.98648586181491e-11 | sushi, nidogen and EGF-like domains 1 |
| ENSMUSG000000055866 | Per2 | 1499.00906731351 | 1.38327282862009 | 0.46808573326219 | 0.111581572373529 | 4.19500929504051 | 2.72860398132914e-05 | 0.0420603493640707 | period circadian clock 2 |
| ENSMUSG000000063415 | Cyp26b1 | 332.600525762669 | 0.466800432924921 | -1.09912219584222 | 0.208426739525057 | -5.27342220267319 | 1.33903030918898e-07 | 0.000652409042394601 | cytochrome P450, family 26, subfamily b, polypeptide 1 |
| ENSMUSG000000079845 | Xlr4a | 21.6024989680628 | 43.3537640434512 | 5.43808535087374 | 1.10971283479549 | 4.90044377280355 | 9.5620421619432e-07 | 0.00310591066156852 | X-linked lymphocyte-regulated 4A |

**Figure 1.**
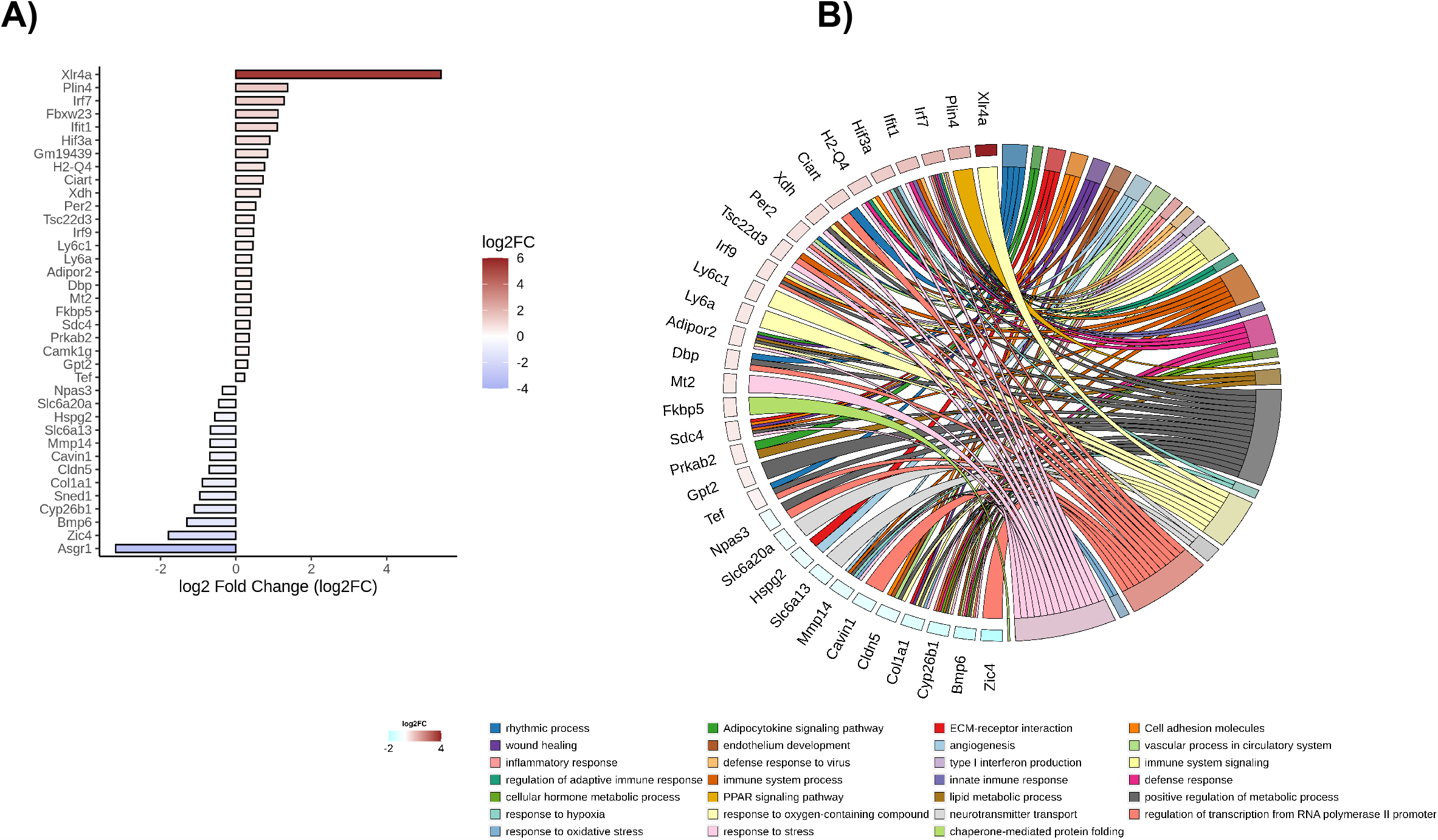
**A**. Log2 Fold Change Bar Plot. The log2 fold change (log2FC) bar plot illustrates gene expression variations of differential expressed genes (including both cortex and hippocampus DEGs). The color scheme assigned blue to downregulated genes and garnet to upregulated genes. Utilizing the ggplot2 library, the log2FC values were plotted on the x-axis, while gene IDs were reordered based on their log2FC values. **B**. GOChord plot showcases the connection between chosen pathways and the genes linked to Type 2 Diabetes Mellitus (T2DM). Genes are depicted in a spectrum of blue-to-garnet colors, representing their log2-Fold-Change values. This visualization offers insight into the relationships between pathways and genes, enhancing our understanding of their potential roles in T2DM.

**Figure 2.**
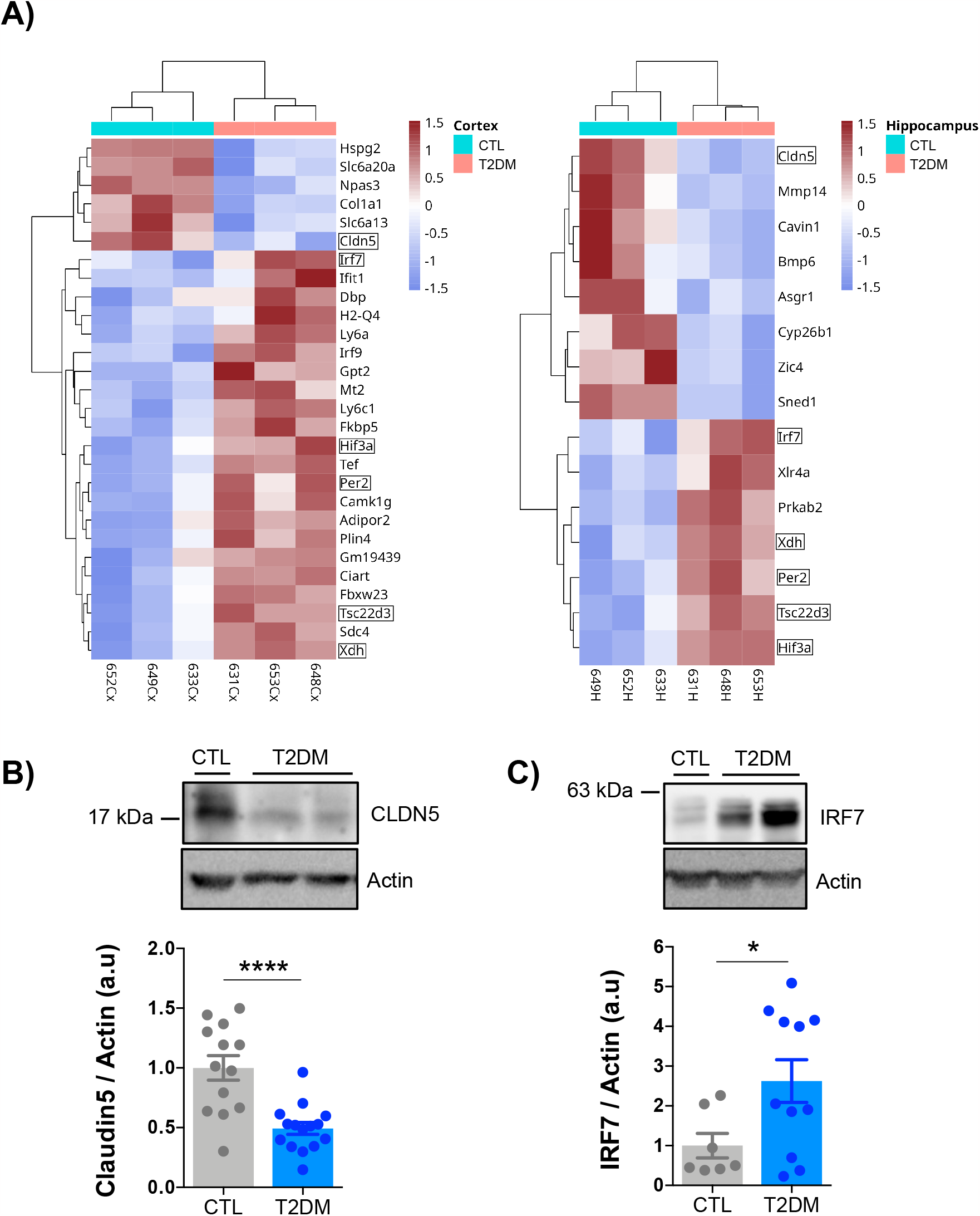
**A. Heatmap** representation of Differential Gene Expression (DEGs) with q-value<0.05 identified in Cortex (left) and Hippocampus (right) between the Type 2 Diabetes Mellitus (T2DM) and control (CTL) WT mice using hierarchical cluster analysis. Each row represents a single gene and each column represents a condition sample. Gene intensities are log2 transformed and displayed as colors ranging from blue to garnet representing the changing process from downregulation to upregulation as shown in the key. **B-C. Immunoblotting** of hippocampal extracts from T2DM and control mice with antibodies against CLDN5 (B, CTL n=13, T2DM n=15, unpaired t test, ****P< 0.0001, data normalized to the CTL group) and IRF7 (C, CTL n=7, T2DM n=11, unpaired t test, *P=0.0385, data normalized to the CTL group). Note that all samples from T2DM mice exhibited reduced CLDN5 levels whereas not all -same-samples had high levels of IRF7.

## 4. Discussion

Our computational data analysis reveals that the alterations induced by T2DM in the cortex and hippocampus of mice align with the anticipated changes commonly observed in how this disease affects other organs in the body. In fact, the six genes miss-regulated in these two structures are associated to alterations in vascular integrity/permeability, inflammation, metabolism, cellular stress, response to hypoxia, cell migration, and transcription [14]. Nevertheless, the observation that alterations occur at the protein level for CLDN5 and IRF7 give to these two molecules a central role in the initiation of functional changes attributed to T2DM.

How could the loss of CLDN5 by T2DM then affect brain function? CLDN5 is a protein that plays a crucial role in the formation and maintenance of tight junctions between endothelial cells in various tissues, including the BBB and blood vessels [9]. Since tight junctions are important for controlling the diffusion of molecules between cells and regulating the permeability of tissues [15] it appears reasonable to claim that the reduced levels of CLDN5 would result in vascular endothelial dysfunction. This in turn would lead to increased vascular permeability, allowing molecules and cells to cross the blood vessel walls more easily triggering, among other consequences, the inflammatory response. This scenario is supported by the increased levels at the mRNA and protein level, of IRF7, a transcription factor involved in the regulation of inflammation, innate immune responses and interferon production [12,16]. While IRF7 is not reported to directly cause cell damage, overactivation of IRF7 could contribute to cell damage and tissue injury through the production of excessive amounts of interferons and other inflammatory molecules. In fact, activation of Interferon Regulatory factor 7 (IRF-7) is responsible for the expression of Type I Interferons and related cytokines. For both humans and mice, the major type I IFN-producing cells in response to viruses, parasites, and bacteria are plasmacytoid dendritic cells (pDCs) in the choroid plexus. In addition, several studies also showed that microglia, oligodendrocytes and neurons can also express type I IFNs. In this regard, it was shown that neurons in areas close to tumor tissue produce IFNß, triggering the production of death-ligand and inducing apoptosis of glioma cells. Early data has shown that chronic activation of type I IFN response in the choroid plexus results in cognitive decline in old mice [17]. It is also worth noting that type I IFNs can be produced by neurons in response to excess amyloid beta-peptide via activation of Toll-like receptors [18]. Furthermore, consistent with the possibility that the loss of CLDN5 by T2DM produces abnormal inflammatory activity in the brain, we have shown the existence of elevated mRNA levels of several other inflammatory intermediaries: *Irf9* (Interferon regulatory factor 9), *Ifit1* (Interferon-induced protein with tetratricopeptide repeats 1), *H2-q4* (Histocompatibility 2, Q region locus 4), *Ly6C1/Ly6a* (Lymphocyte antigen 6 complex) (see Fig. 1A-B and Fig. 2A).

In conclusion, our data strongly indicate that vascular alterations are among the initial effects of T2DM on the brain, achieved through the downregulation of the tight junction protein CLDN5. Further work is required to determine whether CLDN5 decreases as a consequence of the effect of hyperglycaemia on the expression of inhibitory microRNAs [19] or through the activation of inflammatory mechanisms [20]. The fact that all T2DM mice have low levels of CLDN5 in cortex and hippocampus but not all have elevated levels of IRF7 (see Fig. 2 B-C) would be more consistent with the first possibility. In any case, in addition to the mechanistic interpretation, our work opens the possibility of targeting INFs pathway to prevent the onset and progression of cognitive defects in individuals with T2DM.

## Abbreviation list

T2DM: Type 2 Diabetes mellitus
HFD: High fat diet
AD: Alzheimer’s disease
STZ: streptozocin
NBT: nest building test
EPM: Elevated Plus Maze
BBB: Blood brain barrier
DEG: differentially expressed genes
WT: wild type
STZ: streptozotocin
*Hif-3α/*HIF3a: hypoxia inducible factor 3 alpha gene/protein
*Irf7*/IRF7: interferon regulatory factor 7 gene/protein
*Per2/PER2*: period circadian clock 2 gene/protein,
*Xdh*/XDH: xanthine dehydrogenase gene/protein
*Tscd3*/TSCd3: Transforming growth factor β-stimulated clone 22 gene/protein
Cldn5/CLDN5: Claudin-5 gene/protein

## Author’s contributions

CGD designed the overall approach, coordinated the study and drafted the manuscript. MCC and IBL did all biological studies. SGdF carried out the gene expression and pathway analysis. BA supervised all systems biology studies. FXG and EP gave experimental and conceptual supervision and helped prepare the manuscript. All authors read and approved the final manuscript.

## Acknowledgments

The next-generation sequencing (NGS) data analysis has been performed by the Genomics and NGS Core Facility at the Centro de Biología Molecular Severo Ochoa (CBM, CSIC-UAM). The confocal microscopy analysis was performed at the Confocal Microscopy Facility of CBM.

## Data Availability

All data supporting the findings of this study are available from the corresponding authors upon request. The Illumina paired end reads (FASTQ) generated for this study are available at The European Nucleotide Archive (ENA; http://www.ebi.ac.uk/ena/). The RNA raw sequencing data from mouse cortex in control conditions were previously submitted with the study accession number PRJEB61249 and used in another study.

Mouse hippocampus RNA raw sequencing data in control and High fat diet/Steprotozotocin conditions have been deposited under the study accession number PRJEB61724. The dataset pertaining to the High-fat diet/Streptozotocin conditions in the mouse cortex was also submitted under this same study.

