## Supplementary Material for "LOSS OF CLDN5 -AND INCREASE IN IRF7- IN THE HIPPOCAMPUS AND CEREBRAL CORTEX OF DIABETIC MICE AT THE EARLY SYMPTOMATIC STAGE"

**Animals and study design**

All the animal procedures were reviewed and approved by the ethics committees of the Centro de Biología Molecular Severo Ochoa’s (CBM) and Dirección General de Medio Ambiente de la Comunidad Autónoma de Madrid (PROEX 204/19), and performed in accordance with the guidelines of the European Union (2010/63/UE).

Adult male and female WT mice of C57BL/6J background were used in this study. All the animals were housed in the CBM animal facility in standard-sized cages in a ventilated rack, temperature- and humidity-controlled, on a 12-h light cycle (8 AM to 8 PM) with free access to food and water. The mice were randomly divided into two groups with standard rodent chow or a diet containing 60% kcal as fat (HFD) ad libitum. Chronic T2DM was induced by subjecting the mice to a combination of HFD and multiple low doses of STZ, as described in [3]. To minimize confounding effects, whenever possible both control and T2DM groups were handled in parallel (e.g. weight and blood glusose measurements, transfer to behavioral room).

All efforts were made to avoid or minimize pain, distress and discomfort of the experimental animals. Both the animal facility staff and the research team monitored animals weekly. Health was monitored by weight (every two weeks), food and water intake, and general assessment of animal activity, panting, and fur condition. Animals that died prematurely, preventing the collection of behavioral and biochemical data, were not included in the study. Animals that developed spontaneous diseases (e.g. dermatitis, tumor growth) were not included in the study, and, when necessary, subjected to euthanasia.

**Immunoblot**

Cortical and hippocampal homogenates were prepared and subjected to immunoblotting as previously described [3]. In brief, proteins were separated by SDS-PAGE, transferred to nitrocellulose membranes and probed with specific primary antibodies listed below. Bands corresponding to the protein of interest were quantified using FIJI software and normalized with respect to the values obtained for the loading control protein.

**Antibodies**

mouse anti-β-Actin, dilution 1:10000 (Sigma A5441); rabbit anti-Claudin-5, dilution 1:1000 (Cell Signaling 49564); rabbit anti-IRF7, dilution 1:750 (Cell Signaling 72073).

**RNA-sequencing and data analysis**

RNA extraction, sequencing and analysis were performed as we have previously described [3]. In brief, differentially expressed genes (DEGs) between each group were obtained using DESeq2. Genes with *q*-values <0.05 were selected as DEGs. Finally, heatmaps was drawn using the pheatmap R library (version 1.0.12), to perform hierarchical clustering of the DEGs.

To depict gene expression differences, a log2 fold change (log2FC) bar plot of DEGs from both cortex and hippocampus comparatives, was crafted. Using the ggplot2 R package, log2FC values were mapped to the x-axis, and gene IDs were sorted by log2FC magnitude.

**Enrichment Analysis**

An overrepresentation analysis (ORA) [7] using the ClusterProfiler R package [8] was conducted to determine whether known biological functions or processes were over-represented in the experimentally-derived gene list (GO-terms). To enhance the comprehensiveness of the analysis, the recently updated DAVID bioinformatics resource was employed. This allowed for a broader exploration of the data, encompassing searches across various databases such as GO, Reactome and KEGG. The GOChord function from the Goplot R package was implemented to generates a circularly composited overview of selected genes and their assigned functional pathways.

**Statistical analysis**

No statistical methods were used to calculate sample sizes, but our sample sizes are similar to those reported in previous studies [3]. Although experimental conditions were not blinded, data analysis was performed blind whenever possible. The number of individuals used per experimental condition and experimental repeats is indicated in the figure legends. Data are presented as mean ± SEM. In all bar plots, individual data points are presented. Data were analyzed with GraphPad Prism 6. For comparisons between control and T2DM groups, two-tailed unpaired t tests or two-tailed Mann–Whitney tests (when data were not normally distributed) were used. P-values < 0.05 were considered significant. Whenever possible, exact P-values are reported in the figure legends, *n.s* denotes not significant.
